# Multi-stem-loop constrained library enables accelerated SELEX for aptamers with superior kinetics and affinity

**DOI:** 10.1101/2025.11.27.690911

**Authors:** Qinuo Wang, Yuhan Zhao, Siyi Zhao, Anyang Wu, Linfeng Hao, Tianduo Hong, Long Hong

**Affiliations:** School of Life Sciences, Peking University; Beijing, 100871, China; College of Life Sciences, Zhejiang University; Hangzhou, 310058, China; College of Bioscience and Resource Environment, Beijing University of Agriculture; Beijing, 102206, China

## Abstract

Aptamers are single-stranded nucleic acids with ligand-binding capacity as cost-effective alternatives to antibodies. However, their utility is often undermined by the inherent conformational instability of single-stranded nucleic acids, which can compromise binding kinetics and final affinity. To overcome this critical limitation, we synthesized a novel primary library incorporating multiple inlaid stem-loop structures to enhance molecular stability and rigidity. This structural constraint dramatically accelerated the selection process, allowing us to isolate aptamer candidates with superior affinity in just three rounds of SELEX (Systematic Evolution of Ligands by EXponential enrichment), a significant reduction compared to conventional methodologies. Critically, the resulting aptamer demonstrated higher affinity and faster binding kinetics compared to previously reported sequences, which directly validate our hypothesis that structural stabilization leads to high-performance aptamers. This methodology, which integrated rational conformational constraint with a high-throughput selection process, offered a generalizable strategy to efficiently select aptamers with excellent kinetic and thermodynamic performance for demanding applications such as continuous, real-time ligand detection.

## 1. Introduction

Molecular recognition technologies, exemplified by the applications of antibody, have fundamentally transformed both scientific research and clinical medicine. Labeled antibodies are indispensable tools for specific molecule identification (e.g., Western blotting[1]), analysis of molecular interactions (e.g., immunoprecipitation[2–4]), and spatial localization in tissues (e.g., immunohistochemistry[5, 6]) in a plethora of biological researches. Clinically, antibody-based techniques like the Enzyme-Linked Immunosorbent Assay (ELISA)[7–11] are widely used for disease diagnostics, and numerous therapeutic monoclonal antibodies (mAbs), including Rituximab[12–14] and Trastuzumab[15–19], have received FDA approval.

Despite their success, current antibody technologies face several inherent limitations that restrict the ongoing demand for high-affinity and high-specificity molecular recognition elements. First, antibody production is complex and costly. Manufacturing antibodies is reliant on *in vivo* animal or cell culture systems, followed by complex protein purification[20–22], leading to high cost[22–24] and substantial batch-to-batch variability[25, 26]. Second, antibodies suffer from structural instability. As protein molecules, antibodies require stringent storage and handling conditions, making them susceptible to denaturation and resulting in unstable performance.[27] Finally, chemical modification of antibodies is relatively difficult and restricted[28], which limits their functional versatility in diverse research and clinical applications. Consequently, there is a substantial demand for a molecular recognition agent that offers superior stability, lower manufacturing costs, faster development timelines, and simpler functional modification.

Aptamers are a class of *in vitro* synthesized, single-stranded DNA or RNA oligonucleotides capable of binding specifically to a wide variety of ligands.[29, 30] Aptamers targeting a specific molecule are isolated from a vast random nucleic acid library through an iterative selection process known as Systematic Evolution of Ligands by EXponential enrichment (SELEX).[31] Compared to antibodies, aptamers offer numerous advantages, including easier production procedure, facile chemical modification[32], and excellent stability for long-term storage and transportation[29, 33, 34]. These properties endow aptamers with enormous potential in protein qualitative and quantitative analysis, early cancer screening, and pathogen detection.[29, 34, 35]

Despite the distinct advantages of aptamers over antibodies, aptamer-based detection technologies are not yet widely adopted in commercial or clinical settings.[36, 37] A central reason for this limited translation is the inherent flexibility of single-stranded nucleic acid molecules, which compromises the conformational stability of the aptamers and, consequently, their binding performance. In the following section, we will detail the mechanistic reasoning of improving aptamer structural rigidity.

### 1.1. Rationale for enhancing aptamer structural rigidity

For an aptamer to form a stable complex with its target ligand, it must first fold from a highly flexible random coil into a specific, thermodynamically favorable three-dimensional conformation. This necessary folding step introduces the following detriments to binding efficiency.

#### Thermodynamical issue

The binding affinity of an aptamer is quantitively described by the equilibrium constant of the association reaction, which is determined by the standard free energy change: 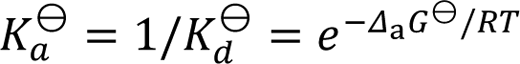. Both binding enthalpy and entropy would affect free energy change: 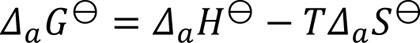. The transition from a highly random state to a specific folded structure involves a large decrease in entropy, which significantly reduces the overall negative Gibbs free energy change of the binding reaction (*Δ_a_G*^⊖^), leading to lower binding affinity (lower *K_a_*^⊖^ and higher *K_d_*^⊖^) compared to an optimally rigid structure.

#### Kinetic issue

The conformational change required for specific binding can act as a rate-limiting step in the overall association process. This results in a slower binding rate (*k_on_*), which negatively impacts the time resolution and sensitivity of aptamer-based biosensors. Such a kinetic compromise would especially limit the application of aptamers in real-time ligand detection, an area of growing interest in both academia and industry.[38–41]

Therefore, engineering aptamers with enhanced structural rigidity is a critical, unmet need for unlocking their full potential in high-performance diagnostic and sensing applications. Analysis based on polymer physics confirms that short double-stranded nucleic acid segments possess significantly enhanced structural rigidity compared to their single-stranded counterparts.[42] This principle is mirrored in nature by highly stable functional nucleic acids, such as tRNA, which utilize stem-loop secondary structures to maintain their defined active conformations.[43] Based on these observations, we hypothesized that artificially constraining the randomized library with multiple stem-loop structures would reduce the unfavorable entropic cost of folding. Although many previous studies have incorporated stem-loops as part of their structured libraries, they either left long unstructured areas[44–46], which could still bring about large entropy decrease upon binding, or based their design on the template of known aptamers to target certain molecules[47] and could not be generalized to a wider range of ligands. Here, we engineered a novel primary SELEX library incorporating multiple inlaid stem-loop structures within the randomized sequence, which provides a stable structural framework while at the same time preserved binding potential for various target molecules. We anticipate that this structural pre-organization accelerates the selection process, enabling rapid identification of aptamers with both high affinity and improved kinetic performance.

## 2. Methods

### 2.1. Oligonucleotide information

All the single-stranded DNAs used in the experiment are listed in Table 1.

**Table 1.**
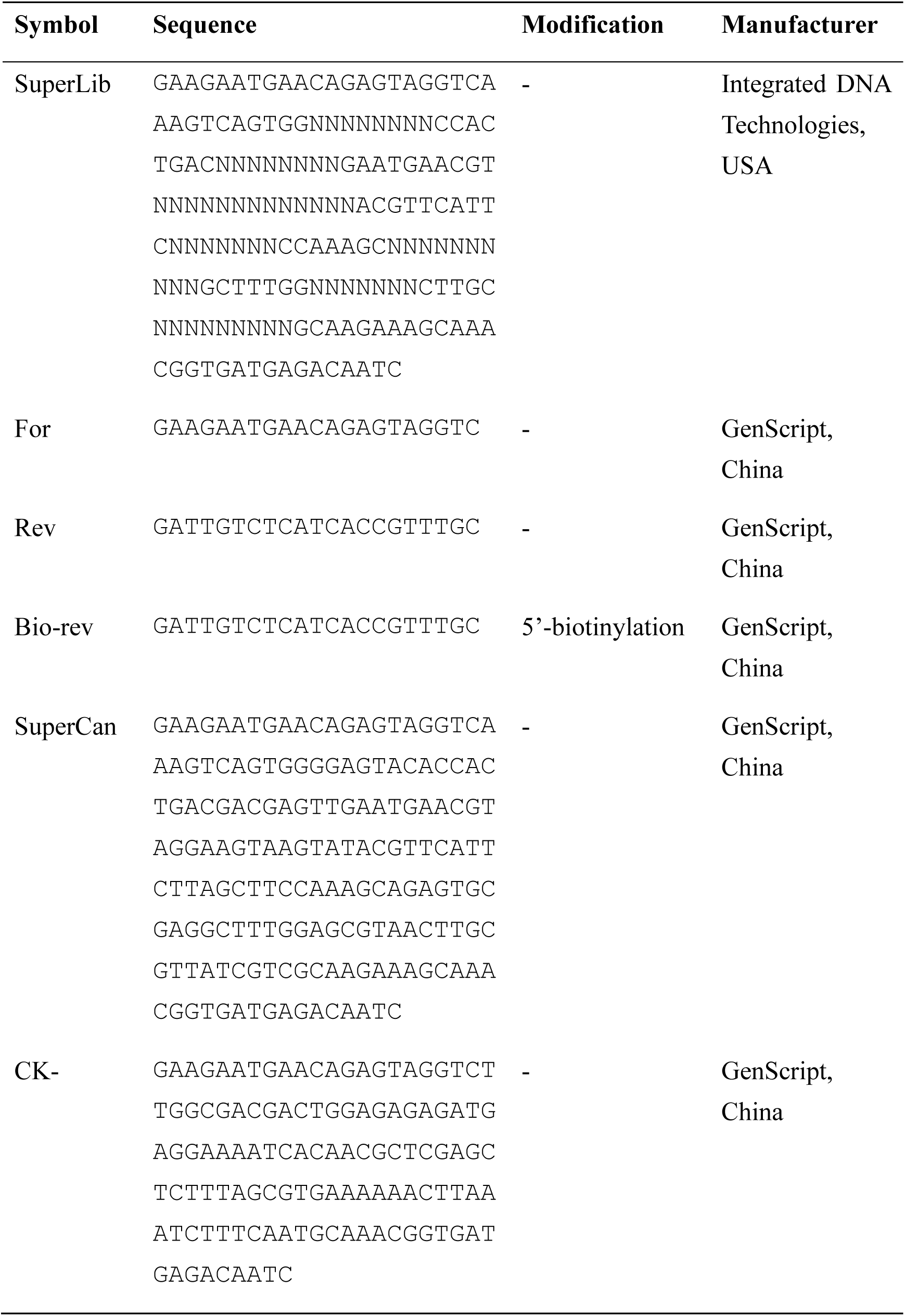
Single-stranded DNAs used in the experiment.

### 2.2. SELEX procedure

#### Enrichment of ligand-binding candidates with Surface Plasmon Resonance (SPR)

SPR was conducted on Biacore T200 (Cytiva, USA). Thrombin (CT11111, Coolaber, China) was coupled onto a CM5 sensor chip (Cytiva, USA) with EDC/NHS chemistry following the manufacturer’s instruction. In each round of enrichment, single-stranded DNA library was dissolved in running buffer (pH 7.4, containing 0.01 M HEPES, 0.15 M NaCl, 3 mM MgCl_2_, 0.005% Tween-20) and passed over the chip surface at a flow rate of 20 µL/min for 2 min. To wash off the oligonucleotides that bound weakly to the ligand, the surface was incubated with the running buffer for 9 times, 30 s each time. Liquid from each time of washing was deposited into 50 μL water and labeled as product 0-8, separately. For product 6-9 in the last round of SELEX, the concentration of Tween-20 in the buffer was raised to 0.05%. After the washing step, the surface was incubated with 0.5% SDS for 30 s to elute the remaining oligonucleotide. Elution liquid was deposited into 50 μL water, ultrafiltered with Amicon® Ultra 30K device (UFC5030BK, Merck, Germany) for 3 times to remove SDS, and labeled as product F. Concentration of the products was measured by qPCR on a StepOnePlus™ instrument (Thermo Fisher, USA), using AceQ qPCR SYBR Green Master Mix (Q111, Vazyme, China).

#### Preparation of the single-stranded DNA library

Single-stranded DNA library was prepared by a novel method combining asymmetric PCR[48] and streptavidin purification. (Fig. 2a) First, the products from SPR enrichment experiment were amplified by asymmetric PCR. The reaction mixture contained 1 × Es Taq MasterMix (Dye) (CW0690, Cwbio, China), 1 µM forward primer (“For” in Table 1), 1/16 µM 5’-biotinylated reverse primer end (“Bio-rev” in Table 1) and the SPR product. The mixture was subjected to 35 cycles at 94°C for 15 s, 56°C for 15 s, and 72°C for 15 s. PCR product was then treated with 1/4 volume of streptavidin magnetic beads (22308, BEAVER, China) for 30 min to remove double-stranded DNA and purified with QIAquick PCR Purification Kit (28104, QIAGEN, Germany). Presence of the product was confirmed by electrophoresis on a 3.5% agarose gel. Purified single-stranded DNA pool was used for next round of SELEX.

#### High-Throughput Sequencing (HTS) of the products

Sequencing library was prepared from product F from each round of SELEX with Hieff NGS® Ultima Pro DNA Library Prep Kit for Illumina® (12201, Yeasen, China) and sequenced on NovaSeq X Plus 25B (Illumina, USA) with PE150 strategy. Raw data was filtered and merged with fastp 0.23.4[49, 50] with default parameters. Merged reads were then counted using a custom-written Python script. Sequence alignment was visualized with ENDscript (https://endscript.ibcp.fr)[51].

### 2.3. Characterization of aptamer candidate

#### Measurement of kinetic parameters with SPR

SPR instrument and sensor chip preparation were as described above. Candidate oligonucleotide of different concentration was dissolved in the running buffer (described above) and passed over the chip surface at a flow rate of 20 µL/min for 3 min, followed by a dissociation period of 5 min. Kinetic parameters were calculated with Biacore T200 Evaluation Software (Cytiva, USA).

#### Validation of binding ability with apta-PCR

Competitive apta-PCR procedure was modified from a previously published method[52]. First, 50 µL of thrombin (20 µg/mL) in a carbonate buffer (50 mM, pH 9.6) was added to each well and incubated for 2 h at room temperature. Following incubation, the wells were washed three times with the wash buffer (pH 7.4, containing 0.01M HEPES, 0.15 M NaCl, 3 mM EDTA, and 0.05% Tween 20). Next, 300 µL of blocking buffer [pH 7.4, containing 0.01M HEPES, 0.15 M NaCl, 3 mM EDTA, 0.05% Tween 20, 1 mg/mL BSA, and 100 µg/mL denatured salmon sperm DNA (R21064, Yuanye, China)] was added to each well and incubated overnight at 4°C. 40 µg/mL of either thrombin or bovine serum albumin (BSA) was mixed with 1 nM of the aptamer molecule in binding buffer (pH 7.4, containing 0.01M HEPES, 0.15 M NaCl, 3 mM MgCl2 and 100 µg/mL denatured salmon sperm DNA). The mixtures were incubated in centrifuge tubes for 2 h at room temperature. Subsequently, 50 µL of the mixture was added to each well of the blocked microplate and incubated for an additional 2 h at room temperature. The wells were then washed three times with the wash buffer. Finally, 100 µL of ultrapure water was added to each well. The plate was incubated at 95°C for 5 min to elute the bound DNA, which was then collected and stored. 2 µL aliquot of the eluted DNA was used for qPCR determination as described above.

#### Calculation of site importance score from HTS data

To identify nucleotide sites important for binding affinity, part of the code from RNA-MobiSeq[53] was used to extract read counts of single-mutated sequences. An importance score defined in Formula 1 was then calculated for each position in the random region of the primary library, in which *I_s_*_,*n*_ is the importance value for position *n* in the candidate sequence *s*, *s_n_* is the base at position *n* in the sequence *s*, *c_s_*_(*n*,*b*)_ is the count value of the sequence in which the base at position *n* in the sequence *s* was mutated to *b*, and *ε* is the pseudo-count (set to 1 throughout the analysis). Secondary structure of the aptamer candidate at 25°C was predicted with RNAStructure 6.4[54] and visualized with pyGenomeTracks 3.9[55, 56]

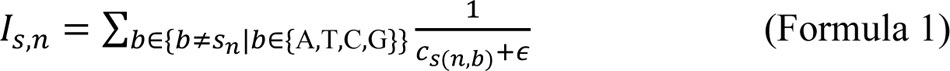

## 3. Results and discussion

### 3.1. Optimization of aptamer primary library with inlaid stem-loops

To enhance the structural rigidity and preserve the versatile binding potential of the primary SELEX library, we introduced four distinct stem-loop structures. These structures featured stem lengths ranging from 5 to 10 bp and loop lengths ranging from 8 to 13 nt, which were separated by random linker sequences (Fig. 1). This arrangement of fixed stems and randomized loops structurally mimics the architecture of an immunoglobulin. In this analogy, the fixed stems serve a function analogous to the Framework Regions (FRs), providing a stable scaffold, while the randomized loops correspond to the Complementarity Determining Regions (CDRs), which primarily determine antigen binding specificity[57]. This novel aptamer primary library was referred to as SuperLib hereafter. We hypothesized that this framework-based design would help stabilize the aptamer’s overall conformation and provide selectable random sites for interaction with various ligands.

**Fig. 1.**
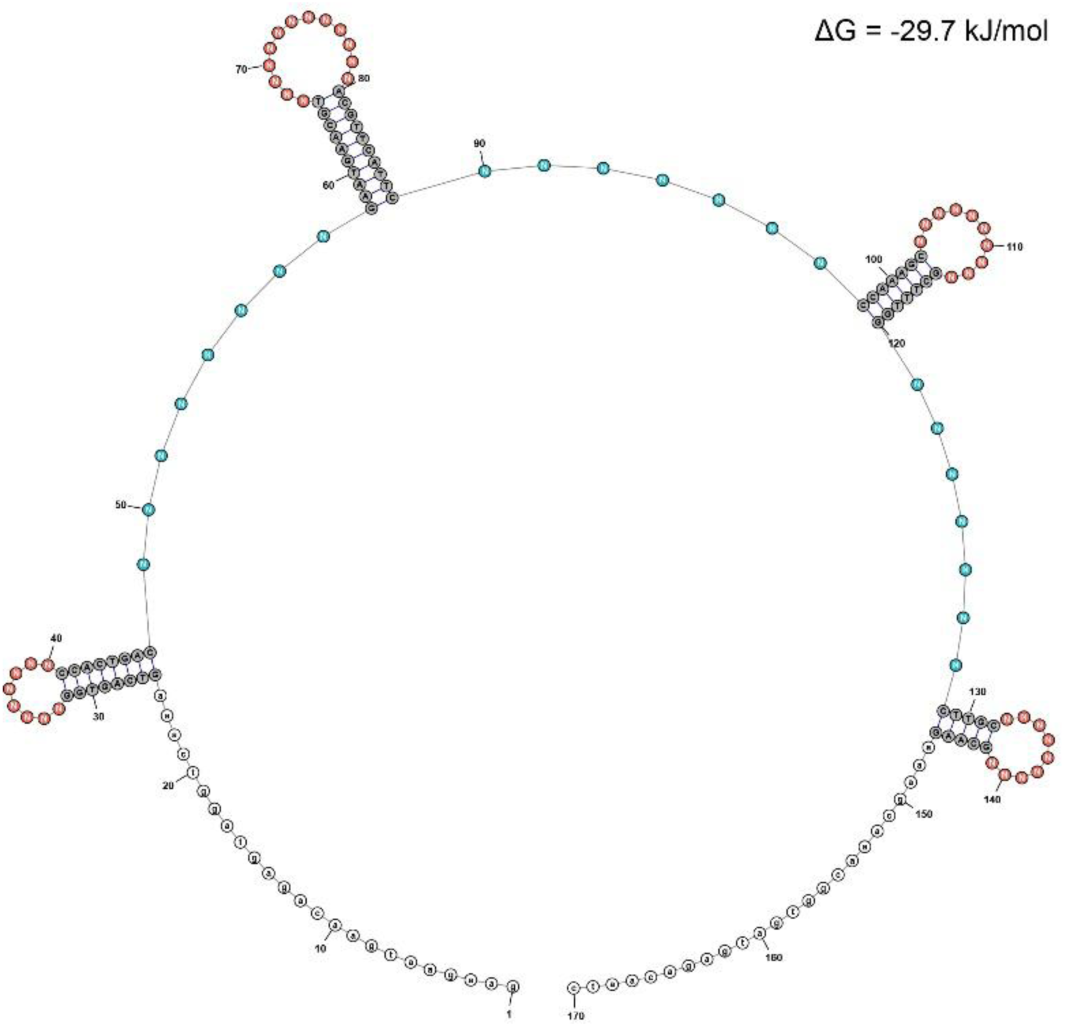
Primary SELEX library inlaid with stem-loops. Predicted folding free energy change Δ*G* for this structure is shown at the upright in this figure. Note that for illustration purpose, pairing in the primer region is ignored; total predicted folding Δ*G* with all these pairings is −31.9 kJ/mol. Red: loops; cyan: linkers; gray: stems; white: primer regions. Both loops and linkers are composed of the random nucleotides (denoted by “N”).

### 3.2. Rapid selection of aptamer candidate from the optimized primary library

For the SELEX procedure, we used SPR for enrichment of ligand-binding candidates and a novel method combining asymmetric PCR[48] and streptavidin purification for preparation of single-stranded DNA (ssDNA) library (Fig. 2). This novel method involves an asymmetric amplification using an excess of the forward primer and a limited amount of 5’-biotinylated reverse primer. The resulting double-stranded DNA (dsDNA) by-product is efficiently cleared using streptavidin magnetic beads. Unlike conventional protocols where labeled dsDNA is captured by streptavidin beads and then denatured to liberate ssDNA[58], our technique minimizes the usage of costly streptavidin beads and bypasses the need for alkaline denaturation, which can compromise the streptavidin-biotin bond and lead to release of interfering amounts of streptavidin and biotinylated DNA[59, 60]. Consequently, this method allows for the high-yield and low-cost production of high-purity ssDNA. (Fig. 2b)

**Fig. 2.**
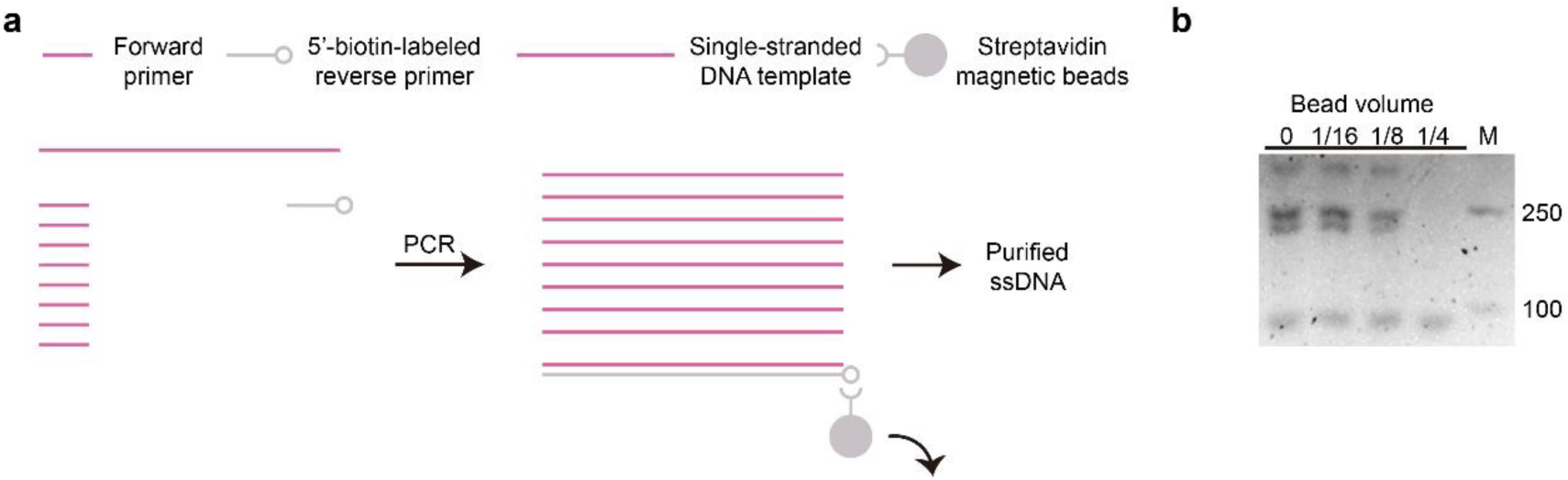
A novel method for single-stranded DNA library preparation. (a) Scheme of the preparation procedure combining asymmetrical PCR and streptavidin purification. (b) Electrophoresis results of the product purified using different volumes of streptavidin magnetic beads. The bead volume is shown as a volume ratio to the PCR product. M: 100 bp Plus DNA Ladder (TransGen, China), with the corresponding double-stranded DNA lengths (in bp) labeled on the right. The single-stranded DNA library length is 170 nt.

In each SELEX round, we quantified the DNA product concentration using qPCR to determine the step-wise recovery rate. The significant increase in recovery rate observed across sequential rounds (Fig. 3a) confirmed the effective enrichment of candidate sequences with target-binding affinity. To increase selection stringency, the Tween-20 concentration was raised during the final four washes of Round 3 (Products 5–8), a change distinctly reflected by the recovery rate curve (asterisk in Fig. 3a).

**Fig. 3.**
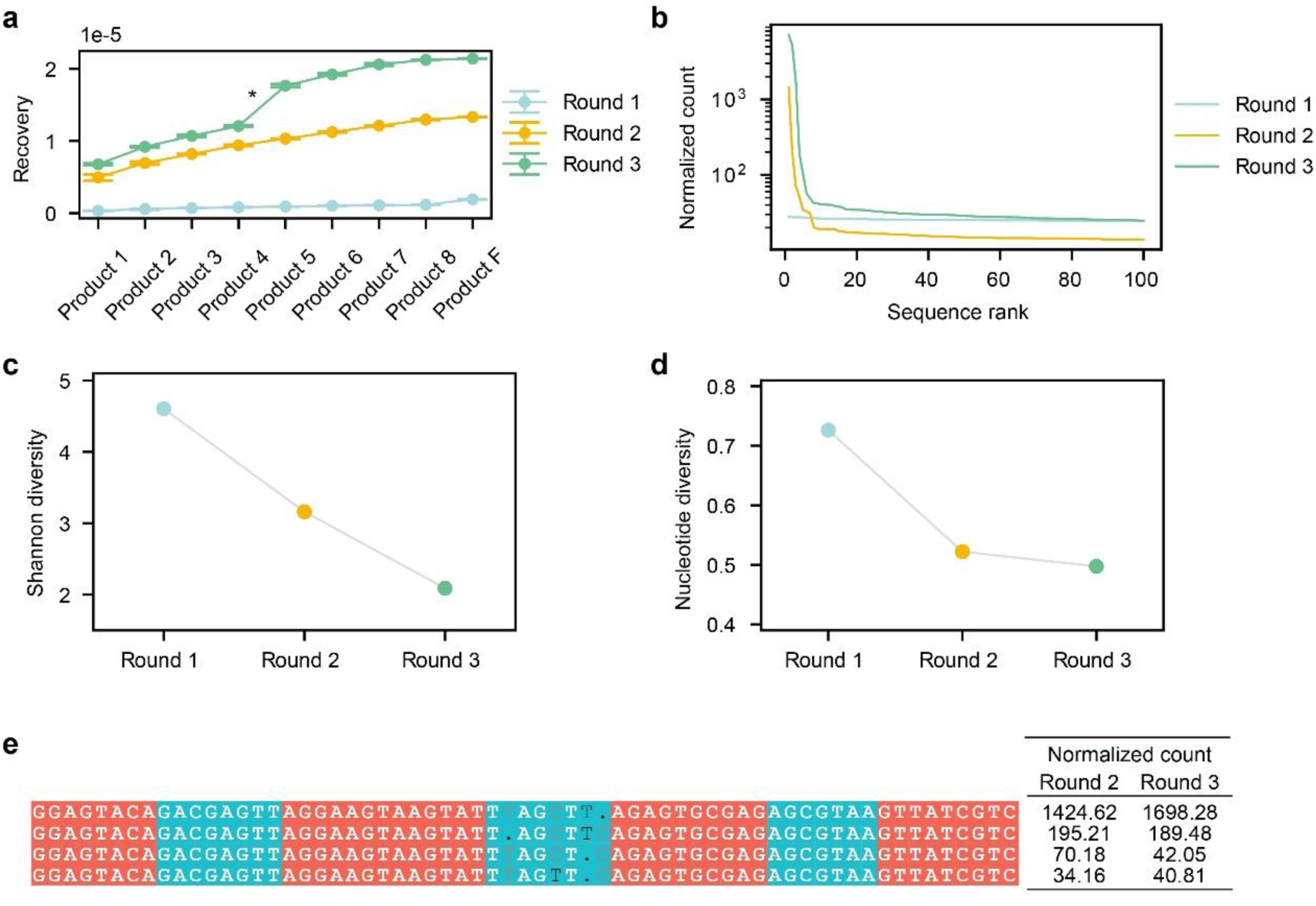
Selection of aptamer candidate through SELEX. (a) Cumulative recovery in each round of SELEX. Asterisk: strengthening of washing in Round 3. (b-d) Top 100 enriched sequences across three SELEX rounds: normalized count per million (CPM), Shannon diversity and average nucleotide diversity. (e) Lead aptamer candidate (first row) and its sequence family. Red: loop regions; cyan: linker regions.

For high-affinity aptamer identification, the eluted pools (Product F) from each round were analyzed via high-throughput sequencing (HTS). The results demonstrated a rapid and effective enrichment of the sequence library. By Round 2, the top-ranked sequence had already gained a 1–2 order of magnitude advantage in read counts over its immediate competitors (Fig. 3b). Successive SELEX round resulted in a persistent increase in the proportion of dominant sequences (Fig. 3b) and a continuous decline in overall sequence diversity (Fig. 3c, d). A consistently high-ranking aptamer candidate (SuperCan hereafter) was identified in the Round 2 and 3 data. This sequence remained in the top 3 across both rounds and was part of a larger, highly similar sequence family, with multiple members consistently ranking in the top 10. This robust enrichment indicated strong binding capacity, leading us to proceed with its synthesis and further characterization.

### 3.3. Characterization of the high-performance aptamer candidate

#### 3.3.1. SPR analysis revealed significantly improved association rate and affinity

We determined the kinetic parameters of SuperCan binding to its ligand using the Surface Plasmon Resonance (SPR) method (Fig. 4a). The results revealed that SuperCan exhibits an extremely high association rate constant (a significantly large *k_on_*) and an extremely high ligand affinity (a greatly reduced *K_d_*). Compared to previously developed conventional aptamers[61–63], SuperCan demonstrated a 1 to 2 orders of magnitude enhancement in its association rate and a 2 to 4 orders of magnitude improvement in ligand affinity (Table 2)[64]. This strongly supports our hypothesis that stabilizing the aptamer’s conformational structure through stem-loop motifs can significantly enhance both its kinetic and thermodynamic performance in ligand binding.

**Fig. 4.**
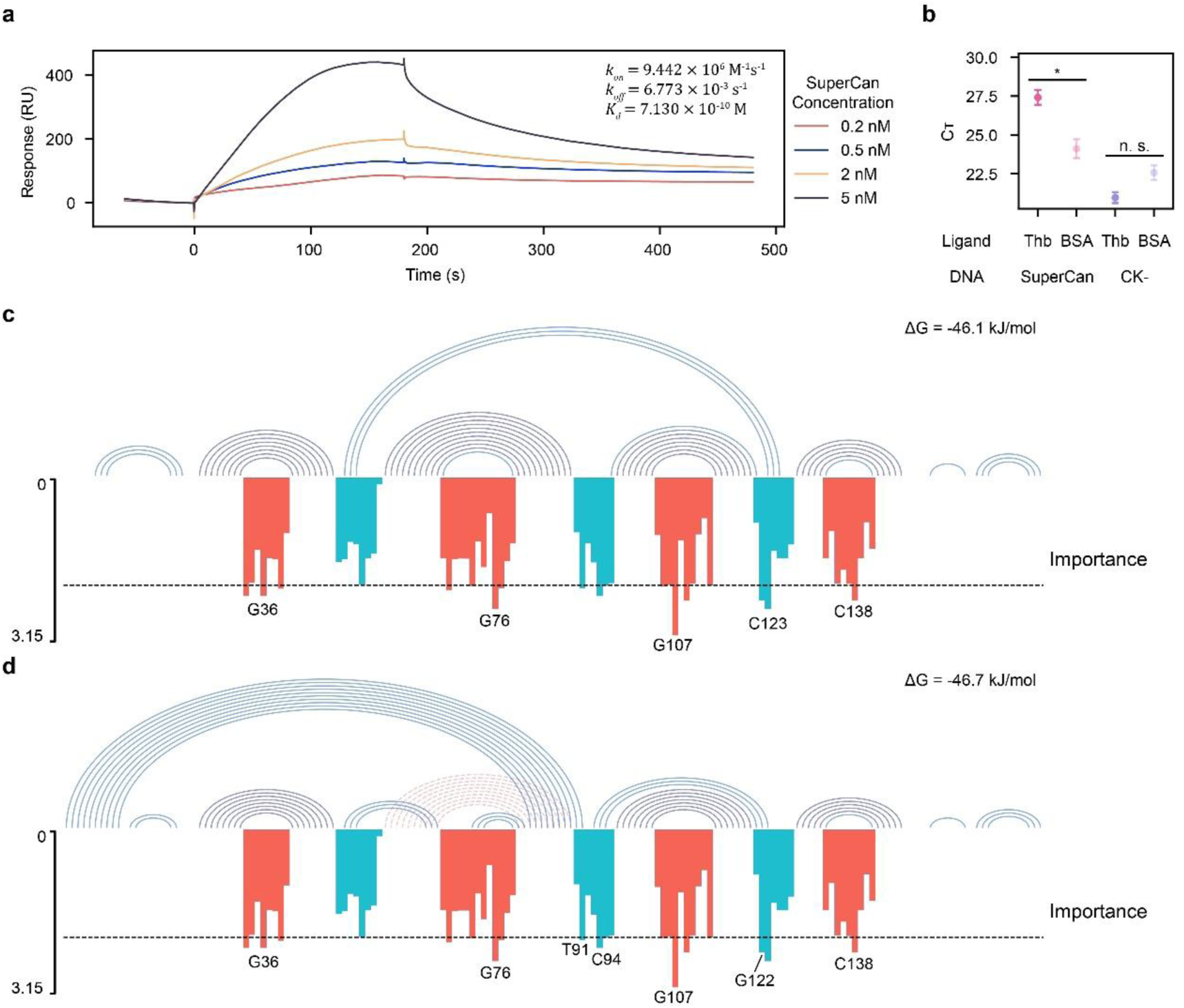
Characterization of the high-performance aptamer candidate. (a) SPR sensorgrams of SuperCan binding to immobilized thrombin. (b) Apta-PCR results of SuperCan binding to thrombin. Thb: thrombin; BSA: bovine serum albumin. Single-tailed t-test was used to test significance. Asterisk: p < 0.05; n. s.: p > 0.05. (c-d) Designed base pairs (red dashed line) and predicted secondary structure (blue solid line, with predicted folding free energy change Δ*G* labeled) of SuperCan, showing the two structures and the importance score for each nucleotide position (black dashed line: 75th percentile). Potential ligand-binding sites and conformation-stabilizing sites are indicated.

**Table 2.** Comparison of binding kinetic parameters with conventional thrombin aptamers.

| Aptamer | $k_{on}$ (M<sup>-1</sup>s<sup>-1</sup>) | $k_{off}$ (s<sup>-1</sup>) | $K_d$ (M) |
| --- | --- | --- | --- |
| SuperCan | $9.442 \times 10^6$ | $6.773 \times 10^{-3}$ | $7.130 \times 10^{-10}$ |
| 15-mer[61, 64] | $3.538 \times 10^5$ | 0.173 | $4.878 \times 10^{-7}$ |
| 29-mer[62, 64] | $8.222 \times 10^5$ | 0.075 | $9.134 \times 10^{-8}$ |
| 32-mer[63, 64] | $1.371 \times 10^5$ | 0.233 | $1.698 \times 10^{-6}$ |

#### 3.3.2. Validation of ligand binding ability with apta-PCR

To further validate the binding ability of SuperCan, we adopted a competitive apta-PCR method[52]. Ligand in the analyte would compete for the aptamer with the immobilized ligand, reducing the amount of aptamer eluted from the plate at last; such a change could be detected by quantitative real-time PCR. A significant increase in the Cт value (i.e., less DNA recovered) was observed with thrombin in the analyte binding to the candidate (Fig. 4b), again illustrating the ligand affinity of SuperCan.

#### 3.3.3. Nucleotide importance revealed potential sites for direct interaction and structural stabilization

To further analyze the distinct roles of individual nucleotide positions in SuperCan during ligand binding, we drew inspiration from the RNA-MobiSeq technique[53] and calculated an importance score for each nucleotide position using the HTS data from the SELEX process. (Formula 1) This approach enabled us to identify several key sites potentially involved in ligand binding within the pre-designed loop structure (G36, G76, G107, and C138 in Fig. 4c-d).

Furthermore, to discover sites that might contribute to stabilizing the aptamer’s conformation, we predicted the secondary structure of SuperCan using RNAStructure[54]. We found that SuperCan formed several new base pairs in addition to the pre-existing stem-loop motif, which significantly enhanced the structural stability of the aptamer molecule (a decrease in the folding Δ*G* of 14.2 kJ/mol). The high importance scores of relevant sites (such as C123 in Fig. 4c) suggest that the conformational stabilizing effect of these new base-pairing structures contributes to the ligand binding process.

Additionally, we observed that SuperCan restructured the pre-introduced base-pairing pattern by forming a new pairing between specific bases in the linker region (T90, T91) and the primer region sequence (A3, A2), resulting in a novel secondary structure pattern (Fig. 4d). This new conformation exhibits a stability similar to that of the conformation in Fig. 4c, suggesting that the two conformations should coexist in equilibrium. The high importance scores of sites closely associated with this new conformation (e.g., T91 and the C94-G122 base pair) imply that this novel conformation may also play a crucial role in ligand binding.

These results indicate that both the sites directly interacting with the ligand and those involved in stabilizing the conformation may have a significant impact on aptamer binding performance, further underscoring the importance of our strategy to stabilize the aptamer’s spatial conformation via the stem-loop structures.

## 4. Conclusion

The application of aptamers in kinetic-demanding fields, such as real-time ligand detection[40, 41], is often limited by the inherent conformational instability of candidates, which negatively affects their binding kinetics (association rate) and affinity. To directly address this challenge, we developed an optimized primary library featuring inlaid stem-loops designed to enhance the structural rigidity of potential aptamer candidates while preserving their binding versatility. This strategic library modification proved highly effective, enabling the rapid selection of a candidate with significantly enhanced fast binding kinetics and high affinity. Ultimately, this work provides a general and effective methodology for accelerating the development of aptamers with superior kinetic performance. We anticipate that this approach will significantly promote the utility of aptamers in various time-sensitive applications, including point-of-care diagnostics and wearable detection systems[38–40].

